# Vigilance profiles at sleep onset in patients with difficulty initiating sleep, with and without depressive or anxiety symptoms

**DOI:** 10.64898/2026.05.05.722943

**Authors:** Laura Abbattista, Benjamin Wacquier, Mélanie Strauss

## Abstract

**Study Objectives:** Difficulty initiating sleep (DIS) is a core insomnia complaint, frequently associated with depressive and anxiety symptoms, yet its objective correlates remain elusive. Insomnia subtypes were merged into a single diagnosis in DSM-5 and ICSD-3 because their criteria lacked reliability, leaving open whether objective markers could help characterize insomnia heterogeneity. Conventional polysomnography (PSG) reduces the wake–sleep transition to a single 30-s epoch. We combined conventional and continuous EEG-based vigilance markers to characterize sleep onset in patients with and without a DIS complaint, and in patients with DIS with and without depressive or anxiety symptoms.

**Methods:** We retrospectively analyzed PSG recordings from 2,952 sleep-clinic patients, grouped by the self-reported complaint of difficulty falling asleep (Insomnia Severity Index item 1 ≥ 2: DIS, n = 1,328; no DIS, n = 1,624). In unmedicated patients with DIS (n = 556), depressive (BDI-II ≥ 14) and anxiety (STAI-T ≥ 45) symptoms were assessed. Vigilance was quantified on 3-s epochs with an intrinsic Vigilance Score (iVS), a probability-of-wakefulness index calibrated on each participant’s own scored night, alongside the theta/alpha ratio and micro-sleep detection, and on 30-s epochs with automated hypnodensities.

**Results:** Patients with DIS showed a longer sleep latency and slightly higher vigilance markers during both wakefulness and sleep, whereas the duration of the SOP itself was only marginally different (median 11.5 min in both groups). Deep sleep also accumulated later across the night (hypnodensity). In wakefulness preceding sleep onset, higher vigilance coexisted with more frequent brief vigilance drops. Among patients with DIS, depressive symptoms were associated with a less wake-like sleep-onset period despite more severe insomnia complaints, a finding not explained by differences in wake amount and convergent with hypnodensity. Anxiety symptoms showed a similar but weaker, non-significant pattern.

**Conclusions:** In patients complaining of difficulty initiating sleep, the delay lies mainly before the first sleep epoch: once it is reached, progression to consolidated sleep is only marginally delayed, although its vigilance content may differ. Continuous EEG markers detect such differences, which are not captured by sleep staging. The association of depressive symptoms with a less vigilant sleep onset, despite worse complaints, suggests that similar insomnia complaints may rest on partly different physiological substrates, and that continuous vigilance markers could help revisit insomnia heterogeneity.

**Statement of Significance:** Insomnia is diagnosed from complaints, and its former subtypes were abandoned because they could not be reliably defined. In nearly 3,000 sleep-clinic patients, we show that the complaint of difficulty falling asleep has objective counterparts: a longer sleep latency and a later build-up of deep sleep. The delay lies mainly before the first sleep epoch, whereas the subsequent transition into consolidated sleep is only marginally prolonged. Continuous EEG markers may further detect subtle differences in the vigilance content of this transition that sleep staging does not capture. Among patients complaining of difficulty falling asleep, those with depressive symptoms showed a less wake-like transition into sleep despite more severe complaints, suggesting that similar complaints may rest on different physiological substrates. Refining these markers could provide an objective dimension to revisit insomnia heterogeneity.

## 1. Introduction

Affecting approximately 10–15% of the general population, insomnia represents a major public health concern due to its detrimental effects on quality of life, mental and physical health (e.g., depression or cardiovascular disease), and society, including increased work absenteeism, disability, and healthcare costs [1]. Since DSM-5 and ICSD-3, insomnia disorder is defined on the basis of complaints of difficulty initiating or maintaining sleep, or early morning awakening, associated with daytime impairment [2,3]. Difficulty initiating sleep (DIS) is thus a symptom rather than a subtype. Although polysomnographic (PSG) studies have frequently reported prolonged sleep onset latency and reduced total sleep time in individuals complaining of DIS [4,5], no robust and clinically useful markers have yet been established.

The sleep-onset period (SOP) is a gradual process during which alertness progressively oscillates toward non-vigilant states, as reflected by the slowing of brain activity from dominant alpha rhythms to theta oscillations [6–8]. Nevertheless, conventional sleep staging defines the sleep onset as the first epoch scored as N1 (or any other sleep stage), thereby reducing the progressive transition into an instantaneous event [9]. This simplification overlooks transient phenomena and rapid fluctuations in vigilance, such as micro-sleep episodes or wake intrusions during sleep, which may be particularly relevant in patients with DIS [10,11]. The scoring of the wake–sleep transition also shows the lowest inter-scorer agreement, and individuals with insomnia tend to overestimate their sleep-onset latency, raising the question of whether conventional PSG lacks the sensitivity to capture subtle increases in vigilance around and during sleep [12–15]. Small case-control studies suggest that patients with sleep-onset complaints show a slower transition from fast-frequency to slower brain activity, with persisting gamma, beta, and alpha rhythms both before sleep onset and during N1 [4,5,16]. Since gamma and beta activity are associated with sensory and cognitive processing, their persistence across the SOP has been interpreted as heightened alertness and cortical hyperarousal, potentially contributing to difficulties initiating sleep [17]. However, these studies included few participants and relied on conventional measures of sleep dynamics, which lack the temporal resolution needed to capture vigilance fluctuations.

Difficulty initiating sleep is frequently associated with depressive and anxiety symptoms, and both major depressive disorder and anxiety disorders are characterized by alterations in sleep continuity and sleep depth [18–20]. The few PSG studies comparing insomnia with and without psychiatric disorders – formerly labelled primary versus secondary or comorbid insomnia – have found that whole-night PSG parameters in major depression with insomnia closely resemble those of insomnia without psychiatric disorder, whereas generalized anxiety disorder with insomnia may be associated with fewer awakenings after sleep onset [21,22]. An earlier study reported higher vigilance with lower delta activity during early N2 sleep in psychophysiological compared with psychiatric insomnia [23], but psychiatric conditions were not specified and only six patients were included per group. To date, PSG studies specifically examining the SOP in relation to depressive and anxiety symptoms remain scarce.

This question has nosological relevance. Earlier classifications distinguished several insomnia subtypes, most of which showed poor reliability and validity. DSM-5 and ICSD-3 therefore merged them into a single insomnia disorder that may co-occur with other conditions [24]. Notably, insomnia related to a mental disorder had been among the most reliably diagnosed categories [25], suggesting that this lumping reflects the lack of reliable criteria for most subtypes rather than evidence of homogeneity. Current efforts to parse insomnia heterogeneity rely either on objective sleep duration, distinguishing insomnia with short versus normal sleep duration with different physiological, cardiometabolic and treatment-response profiles [26], or on self-reported traits and life history [27]. Continuous measures of vigilance at sleep onset could provide an additional objective dimension.

Several EEG-based vigilance markers derived from continuous signal analysis have been proposed. Temporal analysis of the theta/alpha ratio can distinguish between wakefulness (low ratio) and light sleep (high ratio) [28]. Similarly, the odds ratio product (ORP) quantifies the relationship between EEG frequencies over consecutive 3-second epochs and captures variations in sleep depth [29]. By offering finer temporal resolution, these continuous markers allow the characterization of rapid fluctuations in vigilance throughout the SOP. In addition, approaches that estimate the heterogeneity of sleep – by computing the probability of belonging to the different sleep stages at each time-point – such as hypnodensity mapping [30], may provide a more nuanced understanding of insomnia pathophysiology [31].

In this study, we combined conventional PSG measures with continuous EEG vigilance markers – including an individually calibrated index of wakefulness probability – in a large sleep-clinic sample, starting from the complaint rather than from a diagnosis. We asked two questions: i) does the vigilance content of the wake–sleep transition differ in patients complaining of difficulty initiating sleep compared with patients without this complaint; and ii) among patients with DIS, does it differ according to depressive or anxiety symptom burden?

## 2. Materials and methods

### 2.1. Participants

This monocentric retrospective study included patients referred to the Sleep Functional Unit of the Hôpital Universitaire de Bruxelles for an overnight PSG between January 2017 and December 2023 (clinical setting), with approval from the local Ethics Committee (P2024/194). Inclusion criteria consisted of patients who underwent one overnight PSG and completed the Insomnia Severity Index (ISI) questionnaire, corresponding to 3 705 PSG recordings (see flow chart, Figure S1). Recordings from patients with central disorders of hypersomnolence (narcolepsy, idiopathic hypersomnia or Kleine-Levin syndrome) were excluded. Additional recordings were excluded for technical reasons or because the SOP could not be defined according to the criteria described in section 2.4.1. If a patient underwent more than one PSG, only the most recent recording was retained. The final sample consisted of 2 952 recordings and was used to compare vigilance markers between patients with and without a DIS complaint. For the analysis of depressive and anxiety symptoms, additional inclusion criteria required completion of the Beck Depression Inventory (BDI-II) and of the Spielberger State-Trait Anxiety Inventory (STAI) [32,33]. In this second analysis, we excluded patients with active endocrine, inflammatory, or infectious diseases; extreme chronotypes; active substance abuse (including cocaine, cannabis, morphine, alcohol); and current use of hypnotics or psychotropic medications (benzodiazepines, antidepressants, neuroleptics, or opioids). The resulting subsample comprised 556 recordings for the analysis of depressive symptoms and 555 for the analysis of anxiety symptoms.

### 2.2. Questionnaire and clinical information

The DIS complaint was determined using the first item of the ISI, which rates the current severity of difficulty falling asleep from 0 (none) to 4 (very severe) [34]. Patients scoring ≥ 2 (at least moderate difficulty) were classified as having a DIS complaint (DIS, n = 1328), and those scoring < 2 as having no DIS complaint (no DIS, n = 1624). This definition captures a self-reported symptom rather than a diagnosis of insomnia disorder; complaints of sleep maintenance (item 1b) and early morning awakening (item 1c) were not exclusion criteria and are reported in Table 1. Other clinical information was obtained from medical records.

**Table 1.**
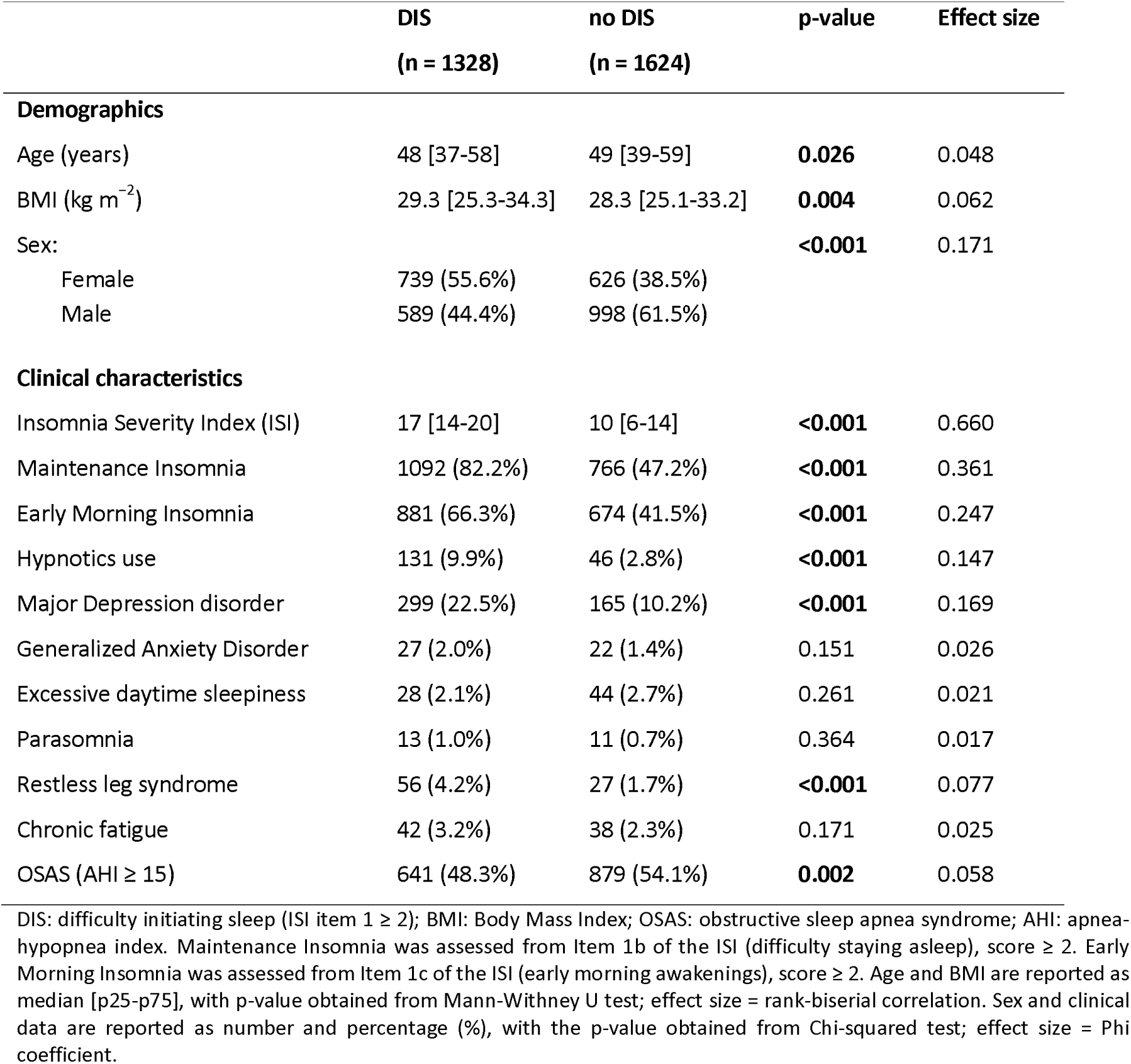
Demographic and clinical characteristics of patients with and without a complaint of difficulty.

|  | DIS<br>(n = 1328) | no DIS<br>(n = 1624) | p-value | Effect size |
| --- | --- | --- | --- | --- |
| **Demographics** |  |  |  |  |
| Age (years) | 48 [37-58] | 49 [39-59] | **0.026** | 0.048 |
| BMI (kg m<sup>-2</sup>) | 29.3 [25.3-34.3] | 28.3 [25.1-33.2] | **0.004** | 0.062 |
| Sex: |  |  | **<0.001** | 0.171 |
| Female | 739 (55.6%) | 626 (38.5%) |  |  |
| Male | 589 (44.4%) | 998 (61.5%) |  |  |
| **Clinical characteristics** |  |  |  |  |
| Insomnia Severity Index (ISI) | 17 [14-20] | 10 [6-14] | **<0.001** | 0.660 |
| Maintenance Insomnia | 1092 (82.2%) | 766 (47.2%) | **<0.001** | 0.361 |
| Early Morning Insomnia | 881 (66.3%) | 674 (41.5%) | **<0.001** | 0.247 |
| Hypnotics use | 131 (9.9%) | 46 (2.8%) | **<0.001** | 0.147 |
| Major Depression disorder | 299 (22.5%) | 165 (10.2%) | **<0.001** | 0.169 |
| Generalized Anxiety Disorder | 27 (2.0%) | 22 (1.4%) | 0.151 | 0.026 |
| Excessive daytime sleepiness | 28 (2.1%) | 44 (2.7%) | 0.261 | 0.021 |
| Parasomnia | 13 (1.0%) | 11 (0.7%) | 0.364 | 0.017 |
| Restless leg syndrome | 56 (4.2%) | 27 (1.7%) | **<0.001** | 0.077 |
| Chronic fatigue | 42 (3.2%) | 38 (2.3%) | 0.171 | 0.025 |
| OSAS (AHI ≥ 15) | 641 (48.3%) | 879 (54.1%) | **0.002** | 0.058 |

DIS: difficulty initiating sleep (ISI item 1 ≥ 2); BMI: Body Mass Index; OSAS: obstructive sleep apnea syndrome; AHI: apnea-hypopnea index. Maintenance Insomnia was assessed from Item 1b of the ISI (difficulty staying asleep), score ≥ 2. Early Morning Insomnia was assessed from Item 1c of the ISI (early morning awakenings), score ≥ 2. Age and BMI are reported as median [p25-p75], with p-value obtained from Mann-Whitney U test; effect size = rank-biserial correlation. Sex and clinical data are reported as number and percentage (%), with the p-value obtained from Chi-squared test; effect size = Phi coefficient.

Among patients with DIS, depressive and anxiety symptoms were assessed with the BDI-II (depressive symptoms: BDI ≥ 14, n = 252; no depressive symptoms: n = 304) and the Trait subscale of the STAI (anxiety symptoms: STAI-T ≥ 45, n = 361; no anxiety symptoms, n = 194) [35,36]. These cut-offs identify clinically relevant symptom burden, not psychiatric diagnoses.

### 2.3. Polysomnographic recordings

The polysomnographic set-up included 2 bipolar electro-oculogram channels, 3 electroencephalogram channels (F4-A1, C4-A1, O2-A1, A1: mastoid reference), 2 submental electromyogram channels, 2 bipolar electrocardiogram channels, a cannula for oro-nasal airflow, a microphone for breath sounds and snoring, thoracic and abdominal inductive belts for respiratory efforts, an oxymeter and 2 pairs of bipolar leg movements channels. In addition, a light sensor was used to record lights-off time and sleep latency (from lights-off to the first sleep epoch). Each recording was visually scored in 30-second epochs by trained technicians according to the American Academy of Sleep Medicine (AASM) criteria [37]. See Figure 1A for an illustration of one night recording.

**Figure 1.**
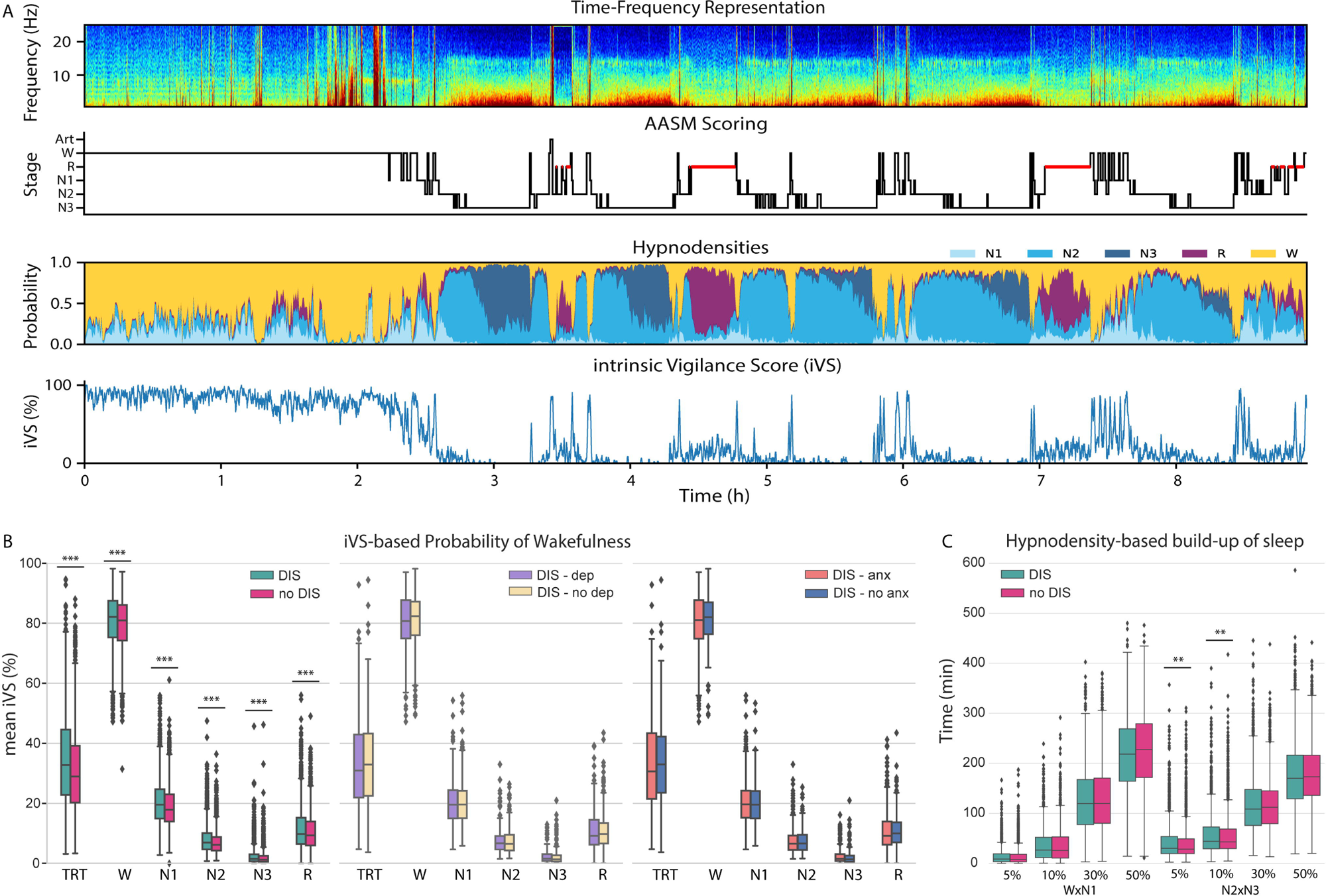
Vigilance analysis across the full night. A. Time-frequency representation, AASM scoring, **Hypnodensities** and iVS across the whole-night recording of one individual. B. Comparison of the mean iVS across the total recording time (TRT) and within each AASM stage between **patients** with and without **DIS** (left), **patients with DIS** with and without depressive symptoms (middle) and **patients with DIS** with and without anxiety symptoms (right). C. Time to reach each threshold of the cumulative sum of the probability products between W and N1 and between N2 and N3 in **patients** with and without **DIS (hypnodensity, independent of individual calibration)**. P-values refer to adjusted logistic regression analyses. * p*_FDR_* < 0.05; p*_FDR_* < 0.01, * p*_FDR_* <0.001.

### 2.4. Sleep Onset Period and vigilance markers

#### 2.4.1. Wakefulness, Sleep Onset Period and Early Consolidated Sleep

The SOP was defined as the interval spanning from the 3 minutes of wakefulness preceding the first epoch of sleep (based on the AASM scoring) to 3 minutes into early consolidated sleep (defined as the first consecutive 10 minutes of N2/N3 sleep, allowing up to 2 min of W or N1). This duration was chosen to capture the physiological transition between the final phase of wakefulness and the initial stage of stable sleep; by construction, the SOP therefore includes 3 min of AASM wakefulness and 3 min of consolidated sleep. The calm wakefulness period before sleep was defined as the 3 minutes wake period starting 6 minutes before the first sleep epoch (used as baseline for continuous vigilance markers).

#### 2.4.2. Theta/Alpha Ratio

To capture rapid fluctuations in vigilance during the SOP (Figure 2A), each recording was segmented into 3-second epochs. consecutive, non-overlapping 3-second epochs. EEG analyses were performed on the C4-A1 derivation. For each epoch, power spectral density (PSD) was estimated with Welch’s method using a 3-s window without overlap, over 0.33– 60 Hz with a frequency resolution of 0.33 Hz, using the MNE-Python software package, v1.8.0 [38]. For each epoch, the theta/alpha ratio was computed as the ratio of the total power within the theta (4-8 Hz) and alpha (8-12 Hz) frequency bands. For each participant, the ratio was normalized by subtracting the mean reference theta/alpha ratio - calculated over the calm wakefulness period - and dividing by its standard deviation (SD). The same theta/alpha ratio reference was used to determine the threshold for microsleep episode (MSE) detection, defined as the mean + 2SD of the reference. A MSE was identified whenever the theta/alpha ratio exceeded this threshold for ≤ 15 seconds. For each event, both its duration and the area under the curve (AUC) above the threshold were computed.

**Figure 2.**
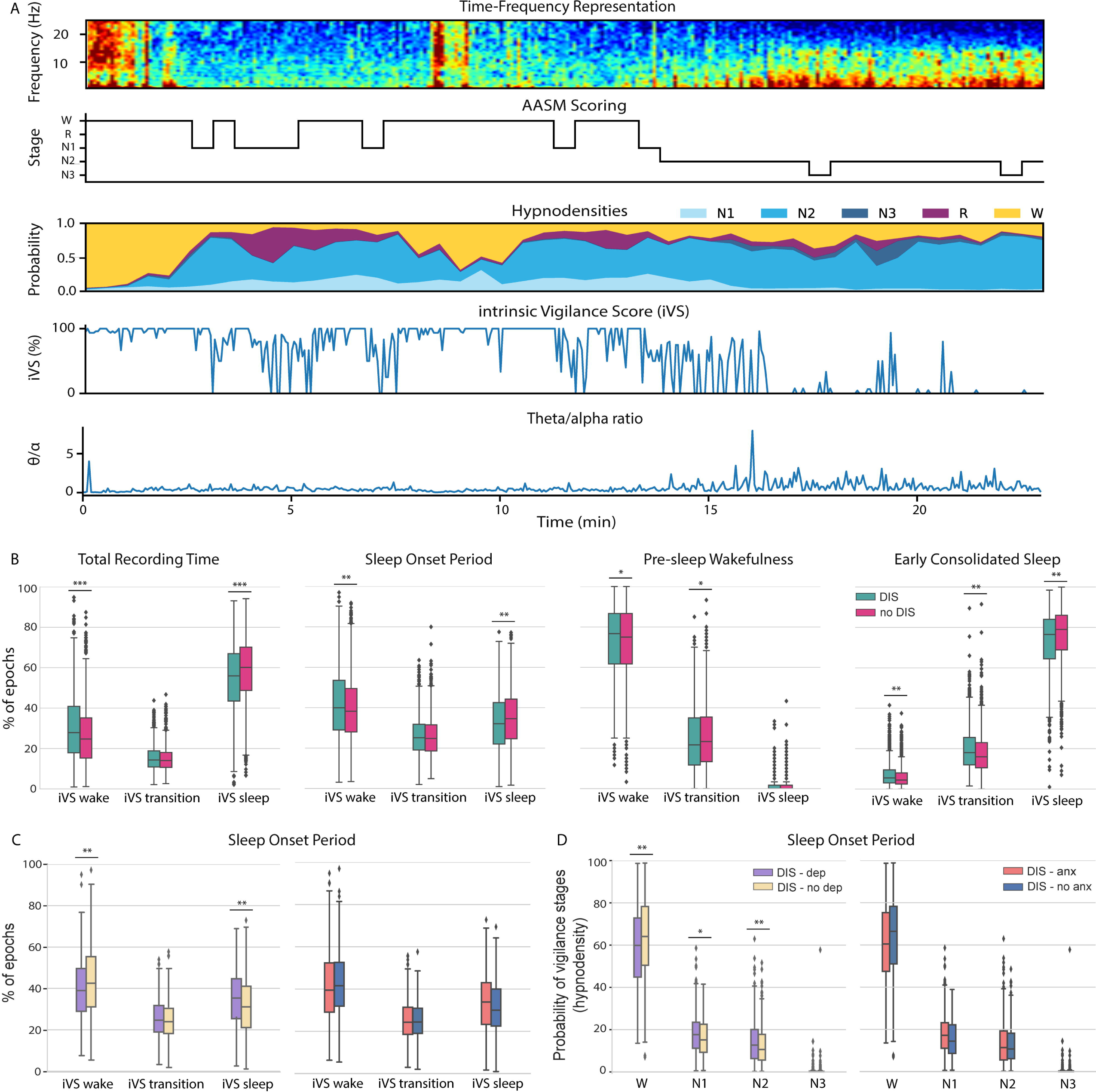
Vigilance analysis across the sleep onset period (SOP). A. Time-frequency representation, AASM scoring, **Hypnodensities**, iVS and the theta-alpha ratio across the SOP and early consolidated sleep of one individual. B. Percentage of 3-s epochs in the different ranges of iVS, classified as sleep (0 – 20), wake (66 – 100) and transition (20 – 66) in **patients** with and without **DIS** across total recording time, SOP, AASM pre-sleep wakefulness and early consolidated sleep. C. Percentage of 3-s epochs in the different ranges of iVS across the SOP, in **patients with DIS** with and without depressive symptoms (left) and with and without anxiety symptoms (right). D. Hypnodensity-derived probability of each vigilance stage across the SOP in **patients with DIS** with and without depressive symptoms and with and without anxiety symptoms. P-values refer to adjusted logistic regression analyses. * p*_FDR_* < 0.05, p *_FDR_* < 0.01, * p*_FDR_* < 0.001.

#### 2.4.3. Intrinsic Vigilance Score (iVS)

To quantify vigilance continuously as the probability of being awake across 3-second epochs, we developed an individualized marker based on the sleep scoring of the participant’s night, the intrinsic Vigilance Score (iVS). For each 3-s epoch, spectral power was summed within four frequency bands corresponding to those used for the odds ratio product (ORP) [29] : delta (0.3–2.3 Hz), theta (2.7–6.3 Hz), alpha (7.3–14 Hz) and beta (14.3–35 Hz). Within each band, power was converted into ten levels defined by deciles within each participant’s night, and each combination of the four levels defined a spectral pattern (bin). Instead of relying on a fixed, group-based lookup table as in the ORP, the probability of wakefulness associated with each bin was estimated from the participant’s own night, as the proportion of epochs in that bin that belonged to AASM-scored wakefulness relative to all epochs in that bin (computed over the whole night, excluding epochs scored as artefact; sleep class: N1, N2, N3 and REM). This individual calibration aims to account for inter-individual differences in absolute spectral power.

The iVS ranged from 0 (0% probability of wakefulness, indicating deep sleep) to 100 (full wakefulness). To determine the iVS thresholds distinguishing three vigilance levels (sleep, transition, wake), we conducted two ROC analyses. In the first one, mean iVS values during consolidated sleep were compared to those during the sleep onset period, while in the second one, values from the sleep onset period were compared to wakefulness. For each ROC curve, the optimal cutoff value was selected as the point that best balanced sensitivity and specificity. This approach allowed to divide iVS values into three categories: < 20 indicating predicted sleep, between 20 and 66 representing the transition state, > 66 indicating wakefulness.

For MSE detection, the threshold was defined as the mean iVS during the calm wakefulness period, minus 2SD. Like the theta/alpha ratio, this threshold captured a stable wakefulness baseline to further detect transient vigilance fluctuations. A MSE was identified whenever the iVS value fell below the threshold for ≤ 15 seconds. The AUC for each MSE was computed as the area between the threshold and iVS curve within the event.

#### 2.4.4. Hypnodensities

The probability of each sleep stage within each 30-second epoch was estimated using the YASA toolbox [39] across the entire recording. We also calculated the cumulative sum of products between the probabilities of specific stages in each epoch and determined the time required to reach 5%, 10%, 30%, 50% of this accumulated sum [30]. These features were computed for two pairs of stages: wake/N1, and N2/N3. The product between stage probabilities reflected the degree of staging uncertainty within each epoch.

### 2.5. Statistical Analysis

Statistical analyses were conducted using Stata version 14.2 and jamovi version 2.6 [40,41]. Data distribution was assessed with the Shapiro-Wilk normality test and visual inspection of histograms. Categorical variables were compared using the Chi-squared test, with effect sizes expressed as Phi coefficients. Continuous variables were compared with the Mann-Whitney U test for non-normal distributions and the independent Student t-test for normal distributions; corresponding effect sizes were calculated with rank biserial correlation and Cohen’s d, respectively.

Associations between the DIS complaint and physiological or clinical markers were assessed using logistic regression models with group (DIS vs no DIS) as the dependent variable and each marker as predictor, adjusting for age, sex, BMI, hypnotic use, major depression, restless leg syndrome, and mixed-obstructive apneas. Odds ratios are therefore expressed per unit of each marker (e.g., per minute or per iVS point). Models comparing patients with DIS with and without depressive or anxiety symptoms included age, sex, BMI, and mixed-obstructive apneas as covariates.

The statistical significance threshold was set at p < 0.05. P-values for sleep-onset markers were corrected for multiple comparisons using the False Discovery Rate (FDR) method. Continuous data are reported as median and percentiles [p25-p75] for non-normal distributions or mean ± SD for normal distributions, and categorical data as n (%).

## 3. Results

### 3.1. Demographic characteristics and clinical information

Patients with DIS were more frequently female, younger, and had a higher BMI compared with those without DIS (Table 1). They also reported greater hypnotics use and a higher prevalence of MDD and restless leg syndrome, but a lower prevalence of obstructive sleep apnea (48.3 vs 54.1%). Most patients with DIS also reported sleep-maintenance difficulties (82.2 vs 47.2%), and patients without DIS had a median ISI score of 10, indicating that the comparison group consisted of sleep-clinic patients rather than good sleepers. Among unmedicated patients with DIS, those with depressive symptoms were more frequently female and reported more severe insomnia (ISI 18 vs 15) than those without (Tables S1a and S1b).

### 3.2. Whole-night sleep

Patients with DIS had a longer sleep latency from lights-off (median 48 vs 41 min), shorter sleep period time (SPT) and total sleep time (TST), lower sleep efficiency, longer latencies to all sleep stages after sleep onset, and increased wake after sleep onset (WASO) (Table 2). DIS was also associated with a reduced proportion of REM sleep – despite fewer microarousals in this stage – and slightly more N2 sleep.

**Table 2.** Sleep and SOP macrostructure variables in patients with and without DIS.

|  | DIS<br>(n = 1328) | no DIS<br>(n = 1624) | OR (95% CI) | p-value |
| --- | --- | --- | --- | --- |
| TIB (min) | 508 [485-539] | 512 [486-539] | 0.999 (0.997-1.000) | 0.066 |
| SPT (min) | 434 [384-475] | 447 [407-479] | 0.997 (0.996-0.998) | **<0.001** |
| TST (min) | 355 [290-408] | 374 [322-417] | 0.996 (0.995-0.997) | **<0.001** |
| Sleep efficiency (%) | 67.7 [55.7-77.9] | 71.5 [61.0-80.4] | 0.978 (0.973-0.983) | **<0.001** |
| N1 (%) | 8.96 [5.65-14.2] | 9.06 [5.87-14.2] | 1.009 (0.999-1.020) | 0.063 |
| N2 (%) | 60.3 [53.5-68.7] | 60.3 [53.6-67.0] | 1.008 (1.001-1.016) | **0.028** |
| N3 (%) | 11.7 [3.54-19.2] | 10.7 [3.75-18.1] | 0.994 (0.986-1.002) | 0.149 |
| REM (%) | 15.7 [10.3-20.2] | 16.7 [12.0-20.2] | 0.980 (0.969-0.991) | **<0.001** |
| Sleep latency (min) | 48 [23.5-93.1] | 41 [19-81] | 1.003 (1.001-1.004) | **<0.001** |
| N2 latency (min) | 3 [1.5-7.5] | 2.5 [1-6] | 1.009 (1.004-1.015) | **<0.001** |
| N3 latency (min) | 31 [18.5-67] | 28 [18-55.3] | 1.003 (1.001-1.004) | **<0.001** |
| REM latency (min) | 113 [74.5-180] | 98 [69.5-152] | 1.001 (1.000-1.002) | **0.013** |
| WASO (min) | 58 [28-106] | 55.5 [29-101] | 1.003 (1.002-1.005) | **<0.001** |
| oAHI (n/h) | 13 [5-27] | 16 [6-33] | 0.997 (0.992-1.001) | 0.169 |
| Desaturation index (n/h) | 9 [2-21.3] | 11 [3-26] | 0.996 (0.992-1.001) | 0.145 |
| μArousal index (n/h) | 13 [8-21] | 14 [8-22] | 0.998 (0.991-1.004) | 0.484 |
| NREM μArousal (n/h) | 11 [6.39-18.5] | 11.8 [6.84-20] | 0.999 (0.992-1.006) | 0.767 |
| REM μArousal (n/h) | 1.57 [0.61-2.99] | 1.90 [0.83-3.46] | 0.955 (0.920-0.991) | **0.015** |
| LPM index (n/h) | 0 [0-5.65] | 0 [0-6.90] | 1.000 (0.995-1.005) | 0.991 |
| SOP length (min) | 11.5 [7-26] | 11.5 [7-23.5] | 1.004 (1.002-1.007) | **<0.001** |
| W duration (min) | 5.5 [3.5-14.5] | 5.5 [3.5-13] | 1.006 (1.003-1.008) | **<0.001** |
| N1 duration (min) | 4 [2-7.5] | 4 [2-7.5] | 1.015 (1.005-1.026) | **0.005** |
| N2 duration (min) | 3 [2-7.5] | 3 [2-7.6] | 1.010 (0.995-1.026) | 0.178 |

TIB: time in bed, SPT: sleep period time, TST: total sleep time, WASO: wake after sleep onset, oAHI: obstructive apnea-hypopnea index; μArousal: micro-arousal; LPM: leg periodic movement. SOP: sleep onset period, from the 3 min of wakefulness preceding the first sleep epoch to 3 min into early consolidated sleep (see Methods). W, N1 and N2 within SOP: cumulative durations of each AASM stage within the SOP. Data are reported as median [p25-p75]. The p-value refers to the logistic regression analysis; OR (95% CI): Odds ratio with 95% confidence interval. p-values for SOP variables are FDR-corrected.

Hypnodensity-derived measures, which rely on a fixed automated model and are therefore independent of individual calibration, showed that patients with DIS required more time to reach 5% and 10% of the cumulative N2×N3 probability product (5%: 30 vs 28.5 min, OR = 1.003 (95% CI, 1.001–1.005), p*_FDR_* < 0.01; 10%: 44 vs 42.8 min, OR = 1.002 (95% CI, 1.001–1.004), p*_FDR_* < 0.05), indicating a delayed build-up of deep sleep across the night. No significant differences were observed for the cumulative W×N1 product (Figure 1C).

The iVS was slightly but consistently higher in patients with DIS across the recording and within each AASM stage (Figure 1B; mean iVS across the total recording time: 32.7 vs 29.0; OR=1.02 (95% CI, 1.02–1.03); W: 82.2 vs 80.9, OR=1.02 (95% CI, 1.01–1.03); N1: 19.5 vs 17.9, OR=1.03 (95% CI, 1.02–1.04); N2: 6.92 vs 6.16, OR=1.06 (95% CI, 1.04–1.08); N3: 1.73 vs 1.42, OR=1.05 (95% CI, 1.02–1.07); REM: 10.1 vs 9.43, OR=1.04 (95% CI, 1.02– 1.05); all p*_FDR_*<0.001). Also, across the full recording, the percentage of epochs classified as wake-like (iVS 66-100) was higher (27.9 vs 24.7%, OR=1.02 (95% CI, 1.01–1.03), p*_FDR_*<0.01), and the percentage of epochs classified as sleep-like (iVS 0–20) lower (55.9 vs 60.1%, OR=0.98 (95% CI, 0.975–0.984), p*_FDR_*<0.01) in patients with DIS (Figure 2B).

Among patients with DIS, the total time in bed (TIB) was slightly higher in those with anxiety symptoms (511 min vs 497 min, OR=1.004 (95% CI, 1.00–1.007), p<0.05), while no other significant differences were found in whole-night macrostructure for either symptom dimension (Tables S2a and S2b). In particular, the amount of wakefulness did not differ between patients with and without depressive symptoms (sleep efficiency 70.1 vs 68.0%, WASO 53 vs 63 min; Table S2a). No significant differences were found between patients with and without depressive or anxiety symptoms in mean iVS across sleep stages (Figure 1B) or in the cumulative W×N1 and N2×N3 products (Tables S3a and S3b).

### 3.3. Sleep Onset Period

The SOP had the same median duration in both groups (11.5 minutes, Table 2), but its distribution extended toward longer values in patients with DIS, resulting in a small but significant difference in adjusted models (DIS = 11.5 [7–26] min, no DIS = 11.5 [7-23.5] min, OR = 1.004 per min (95% CI, 1.002–1.007), p*_FDR_* < 0.001). This difference was carried by slightly longer durations of wakefulness and N1 within the SOP, which, despite identical medians, were longer in DIS in adjusted models (W: OR = 1.006 (95% CI, 1.003–1.008), p*_FDR_* < 0.001; N1: OR = 1.015 (95% CI, 1.005–1.026), p*_FDR_* < 0.01), whereas N2 duration did not differ.

During the SOP, the mean iVS was higher in the DIS group (47.5 vs 45.5, OR=1.01 (95% CI, 1.006–1.017), p*_FDR_*<0.01), with a higher percentage of wake-like epochs (iVS 66-100, 40.0 vs 38.2%, OR=1.01 (95% CI, 1.005–1.014), p*_FDR_*<0.01) and a lower percentage of sleep-like epochs (iVS 0-20, 32.1 vs 34.5%, OR=0.99 (95% CI, 0.983–0.994), p*_FDR_*<0.01, Figure 2B).

MSEs, as assessed with both the theta/alpha ratio and iVS methods, were more numerous and of longer cumulative duration in the DIS group, but their rate per minute did not differ (Tables S4 and S5), indicating that this difference mainly reflected the slightly longer SOP. In contrast with the iVS, the hypnodensity-derived probabilities of wakefulness and sleep stages across the SOP did not differ between groups (Table S6).

SOP duration did not differ between patients with DIS with and without depressive or anxiety symptoms (Tables S7a and S7b). Patients with depressive symptoms showed a less wake-like SOP than those without (mean iVS 45.9 vs 48.4; OR=0.98 (95% CI, 0.97–0.99), p*_FDR_*<0.01), with fewer wake-like epochs (iVS 66-100, 38.6 vs 42.4%, OR=0.98 (95% CI, 0.97–0.99), p*_FDR_*<0.01) and more sleep-like epochs (iVS 0-20, 35.3 vs 30.8%, OR=1.02 (95% CI, 1.01– 1.04), p*_FDR_*<0.01, Figure 2C). This difference occurred in the absence of differences in the amount of wakefulness (Table S2a) and converged with hypnodensity, which is independent of individual calibration: patients with depressive symptoms had a lower probability of wakefulness (59.9% vs 64.1%; OR=0.99 (95% CI, 0.98–0.996); p*_FDR_*<0.01) and higher probabilities of N1 (17.7% vs 15.2%; OR=1.02 (95% CI, 1.003–1.04); p*_FDR_*<0.05) and N2 (12.7% vs 10.6%; OR=1.03 (95% CI, 1.01–1.04), p*_FDR_*<0.01, Figure 2D). This less wake-like SOP was observed although patients with depressive symptoms reported more maintenance and early morning insomnia symptoms (ISI 18 vs 15; Table S1a). Anxiety symptoms were associated with a numerically similar but weaker pattern that did not reach significance after adjustment (mean iVS 46.2 vs 49.3; OR=0.991, p*_FDR_*=0.58; Table S9b). No differences were found in MSEs detected with the iVS or the theta/alpha ratio for either symptom dimension (Tables S8 and S9).

### 3.4. AASM Pre-sleep Wakefulness

During the 3 min of wakefulness immediately preceding the first AASM sleep stage – a window of fixed duration that contains only AASM-scored wakefulness and is therefore not affected by differences in SOP length – the mean iVS was slightly higher in the DIS group (80.5 vs 79.7; OR=1.009 (95% CI, 1.002–1.02), p*_FDR_*<0.05; Figure 2B), along with a higher percentage of epochs of high wakefulness probability (iVS 66-100, 76.7 vs 75.0%; OR=1.005 (95% CI, 1.00–1.009), p*_FDR_*<0.05) and a lower percentage of transitional epochs (iVS 20-66, 21.7 vs 23.3%; OR=0.995 (95% CI, 0.990–1.00), p*_FDR_*<0.05). Despite these signs of higher vigilance, MSEs detected with the iVS were more numerous, more frequent per minute (1.33 vs 1.00 /min), longer and larger in the DIS group (Table S10), whereas no difference was found with the theta/alpha ratio (Table S11).

Depressive or anxiety symptoms were not associated with any significant difference in iVS or theta/alpha ratio measures during pre-sleep wakefulness (Tables S12 and S13).

### 3.5. Early consolidated sleep

The DIS group still showed a higher mean iVS in the early consolidated sleep period (11.6 vs 10.4, OR=1.02 (95% CI, 1.01–1.04), p*_FDR_*<0.01), with more epochs with high vigilance scores (iVS 66-100, 5.5 vs 4.5%; OR=1.03 (95% CI, 1.02–1.05), p*_FDR_*<0.01), but here also more epochs of intermediate sleep (iVS 20-66, 18.0 vs 16.0%; OR=1.015 (95% CI, 1.008– 1.023), p*_FDR_*<0.01), and fewer epochs classified as sleep (iVS 0-20, 76.5 vs 79%; OR=0.987 (95% CI, 0.982–0.993), p*_FDR_*<0.01, Figure 2B). The characteristics of MSEs across this period did not differ between the two groups despite a higher iVS variance (Tables S14 and S15).

Depressive or anxiety symptoms were not associated with significant differences during early consolidated sleep (Tables S16 and S17).

## 4. Discussion

In this large sleep-clinic sample, we examined the wake–sleep transition starting from the complaint of difficulty initiating sleep rather than from an insomnia diagnosis. First, we found that the DIS complaint had objective counterparts that do not depend on individual calibration: a longer sleep latency and a delayed accumulation of deep sleep across the night, whereas the transition from the first sleep epoch to consolidated sleep was only marginally longer. Second, the individually calibrated iVS suggested a slightly more wake-like EEG in patients with DIS during the SOP, in the last minutes of wakefulness and within each AASM stage. Third, among unmedicated patients with DIS, depressive symptoms were associated with a less wake-like sleep onset despite more severe complaints during the night.

The longer sleep latency and the later build-up of deep sleep indicate that, in this population, the complaint of difficulty initiating sleep is not only a matter of perception. The iVS results further suggest that, at equal AASM stage, the EEG of patients with DIS lies slightly closer to wakefulness, from pre-sleep wakefulness to N2, N3 and REM sleep. Such a pattern is consistent with models of cortical and cognitive hyperarousal in insomnia [42–44] and with EEG evidence of elevated high-frequency activity across NREM and REM sleep and of 24-h cortical hyperarousal in insomnia disorder [45,46].

Considering the SOP itself, its median duration was identical in both groups (11.5 min), and the small difference observed in adjusted models was driven by the upper tail of the distribution. Once the first sleep epoch occurs, the time needed to reach consolidated sleep therefore appears largely preserved, and differences rather concern the content of the transition, with higher vigilance states (AASM wakefulness and iVS scores). Also, in the fixed 3-min period of AASM wakefulness preceding sleep onset, patients with DIS showed slightly higher mean vigilance but also more frequent brief vigilance drops, a pattern detected by the iVS but not by the theta/alpha ratio. Rather than simple hypervigilance, this suggests an unstable vigilance system oscillating between wake-like and sleep-like states before sleep onset. In light of experimental and theoretical models describing sleep onset as a bistable or metastable process [8,47,48], this pattern may reflect a less stable control of arousal at the wake–sleep boundary. Across the whole SOP, by contrast, MSE rates did not differ, and the greater number of MSEs mainly reflected the slightly longer SOP.

The analysis of depressive symptoms provides an interesting contribution of our iVS findings. Among unmedicated patients with DIS – thereby excluding direct effects of psychotropic medications – those with depressive symptoms showed a less wake-like SOP than those without, although they reported more severe insomnia. This difference converged with hypnodensity estimates derived from a fixed model. Depression and anxiety are strongly associated with insomnia at the clinical level [49,50], but these results suggest that, for a similar or even more severe complaint, the physiological substrate of the transition into sleep may differ according to depressive symptom burden. Insomnia with prominent depressive symptoms could involve broader alterations in homeostatic, circadian, affective and cognitive processes rather than heightened vigilance at sleep onset [51–53]. In line with this view, major depression is associated with hypersomnolence rather than hyperarousal in a substantial subset of patients [54,55]. Anxiety symptoms showed a numerically similar but weaker pattern.

These findings can be related to current phenotyping approaches. In insomnia with objective short sleep duration, physiological hyperarousal (e.g., autonomic and hypothalamic– pituitary–adrenal activation) has been associated with a depressive-somatic rather than anxious-ruminative profile [24,26]. Our observation of a less wake-like EEG at sleep onset in patients with depressive symptoms is not necessarily at odds with this view, since cortical EEG vigilance and systemic physiological arousal may dissociate; testing whether the present EEG differences vary with objective sleep duration is a direct next step. More generally, insomnia subtypes were merged in DSM-5 and ICSD-3 because their clinical criteria were not reliable, not because subtypes were shown to be physiologically homogeneous [24,25]. Continuous vigilance markers may provide an objective dimension to revisit this heterogeneity, complementary to objective sleep duration and to self-reported trait profiles [27]. Whether they identify stable subgroups with distinct course or treatment response remains to be established.

The iVS was designed to express vigilance as a probability of wakefulness on a 0–100 scale, calibrated on each participant’s own sleep–wake distribution to account for inter-individual differences in absolute spectral power, and integrating information across the delta to beta bands, whereas single spectral ratios such as the theta/alpha ratio are mainly informative around the wake–sleep boundary.

Several limitations should be noted. First, DIS was defined from a single ISI item reflecting a self-reported complaint, not from a diagnosis of insomnia disorder, and most patients with DIS also reported sleep-maintenance difficulties; the present results therefore concern a symptom dimension rather than a clinical subtype, whose temporal stability is known to be limited [56,57]. Second, effect sizes were small, and their clinical significance remains to be established. Third, depressive and anxiety symptoms were assessed with questionnaire cut-offs rather than structured interviews, and the BDI-II includes sleep and fatigue items; the use of dimensional measures nevertheless aligns with contemporary transdiagnostic frameworks, and the exclusion of patients receiving psychotropic medications allowed a cleaner assessment of symptom burden. Finally, only one in-laboratory night was analyzed, and no information was available on perceived sleep latency on the night of the recording. Future studies should test the longitudinal stability of these markers, their relationship with subjective sleep perception and daytime functioning, and their modulation by treatment.

Altogether, starting from the complaint of difficulty initiating sleep, this study shows that the complaint has objective counterparts in sleep latency and in the build-up of deep sleep, and that continuous EEG markers can detect differences in the vigilance content of the wake– sleep transition that are not captured by sleep staging. In these patients, the delay lies mainly before the first sleep epoch, where vigilance showed more frequent brief drops, possibly reflecting unsuccessful attempts to fall asleep; once the first sleep epoch is reached, progression to consolidated sleep is only marginally delayed, although its vigilance content may differ. Also, among patients with DIS, depressive symptoms were associated with a less wake-like sleep onset despite more severe complaints, suggesting that similar complaints may rest on partly different physiological substrates. Testing the stability of individual vigilance markers, their relation to objective sleep-duration phenotypes and their predictive value for treatment response will determine whether they can contribute to a biologically informed description of insomnia heterogeneity, particularly as home-based EEG enables multi-night assessment.

## Acknowledgements

We gratefully acknowledge the support of the Sleep Functional Unit team members and thank all patients who participated in this study for their valuable contribution.

## Funding

M.S. was supported by the Fonds de la Recherche Scientifique – FNRS.

## CrediT

Conceptualization : M.S., L.A., B.W., Methodology : M.S., L.A., B.W., Software : L.A., B.W, Formal analysis : L.A., B.W, Investigation : L.A., Resources : M.S., Data Curation : L.A., B. W., Writing - Review & Editing : M.S., L.A., B.W., Visualization : M.S., L.A., B.W, Supervision : M.S., Project administration : M.S., Funding acquisition : M.S.

## Data availability

data are availabile upon reasonable request.

## Disclosure Statement

### Financial Disclosure

none.

### Non-financial Disclosure

none.

### Preprint repositories

https://doi.org/10.64898/2026.05.05.722943

## Notes

### Competing Interest Statement

The authors have declared no competing interest.

### Summary of Updates

This revised version is based on the same data and analyses as version 1. The main changes are: (1) a new title and a revised framing, starting from the self-reported complaint of difficulty initiating sleep (DIS) rather than sleep-onset insomnia to avoid ambiguity between symptoms and diagnosis; (2) an Introduction and Discussion placing the findings in the context of insomnia nosology and current phenotyping approaches, and update of the references; (3) a more detailed description of the EEG processing and of the intrinsic Vigilance Score (iVS); (4) correction of typographical and numerical errors

