## Supplementary Information for "Vigilance profiles at sleep onset in patients with difficulty initiating sleep, with and without depressive or anxiety symptoms"

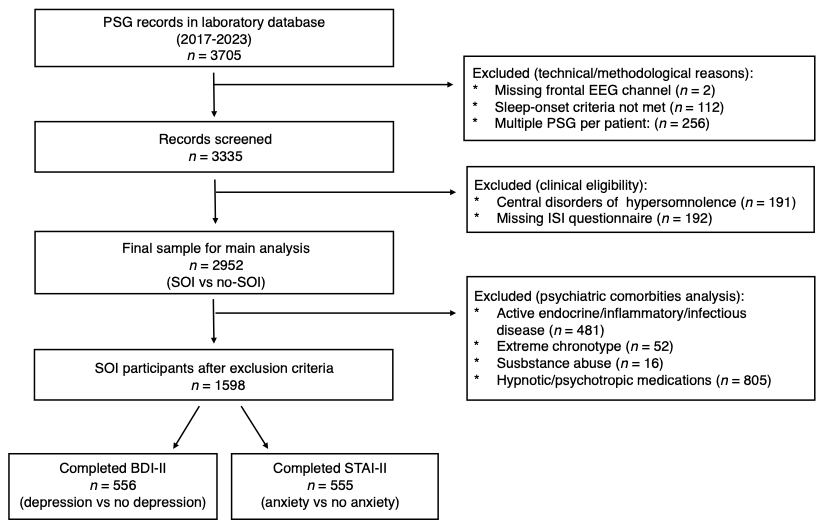

***Figure S1 –*** *Flow Chart.*

Table S1a – Demographic and clinical characteristics in DIS with/without depressive symptoms

| Characteristics | DIS – dep  (n = 252) | DIS – no dep  (n =304) | p-value | Effect size |
| --- | --- | --- | --- | --- |
| Demographics |  |  |  |  |
| Age (years) | 44 [31-55] | 46 [35-58] | 0.051 | 0.0962 |
| BMI (kg m^−2^) | 30.9 [26.0-35.2] | 29.0 [25.6-34.2] | 0.080 | 0.0862 |
| Sex: |  |  | **<0.001** | 0.179 |
| Female | 143 (56.7%) | 118 (38.8%) |  |  |
| Male | 109 (43.3%) | 186 (61.2%) |  |  |
| Clinical characteristics |  |  |  |  |
| Insomnia Severity Index score | 18 [15-21] | 15 [12-18] | **<0.001** | 0.380 |
| Maintenance Insomnia | 219 (86.9%) | 242 (79.6%) | **0.023** | 0.0965 |
| Early Morning Insomnia | 178 (70.6%) | 178 (58.6%) | **0.003** | 0.125 |
| Excessive daytime sleepiness | 9 (3.6%) | 7 (2.3%) | 0.365 | 0.0602 |
| Parasomnia | 3 (1.2%) | 6 (2.0%) | 0.364 | 0.0167 |
| Restless leg syndrome | 5 (2.0%) | 12 (3.9%) | 0.181 | 0.0568 |
| Chronic fatigue | 11 (4.4%) | 9 (3.0%) | 0.366 | 0.0601 |
| OSAS (AHI >=15) | 108 (42.9%) | 160 (52.6%) | **0.022** | 0.0974 |

BMI: Body Mass Index; OSAS: obstructive sleep apnea syndrome; AHI: apnea-hypopnea index. Age and BMI are reported as median [p25-p75], with p-value obtained from Mann-Whitney U test; effect size = rank-biserial correlation. Sex and clinical data are reported as number and percentage (%), with the p-value obtained from Chi-squared test; effect size = Phi coefficient.

Table S1b – Demographic and clinical characteristics in DIS with/without anxiety symptoms

| Characteristics | DIS – anx  (n = 361) | DIS – no anx  (n =194) | p-value | Effect size |
| --- | --- | --- | --- | --- |
| Demographics |  |  |  |  |
| Age (years) | 43 [31-54] | 49.5 [38.3-60] | **<0.001** | 0.207 |
| BMI (kg m^−2^) | 29.2 [25.2-34.3] | 30.9 [26.7-35.3] | **0.023** | 0.117 |
| Sex: |  |  | 0.351 | 0.0396 |
| Female | 175 (48.5%) | 86 (44.3%) |  |  |
| Male | 186 (51.5%) | 108 (55.7%) |  |  |
| Clinical characteristics |  |  |  |  |
| Insomnia Severity Index score | 17 [15-21] | 15 [11-17] | **<0.001** | 0.375 |
| Maintenance Insomnia | 300 (83.1%) | 160 (82.5%) | 0.851 | 0.00795 |
| Early Morning Insomnia | 251 (69.5%) | 100 (51.5%) | **<0.001** | 0.178 |
| Excessive daytime sleepiness | 11 (3%) | 4 (2.1%) | 0.603 | 0.0427 |
| Parasomnia | 6 (1.7%) | 3 (1.5%) | 0.915 | 0.0045 |
| Restless leg syndrome | 12 (3.3%) | 5 (2.6%) | 0.626 | 0.0207 |
| Chronic fatigue | 14 (3.9%) | 5 (2.6%) | 0.551 | 0.0464 |
| OSAS (AHI>=15) | 152 (42.1%) | 113 (58.2%) | **<0.001** | 0.154 |

BMI: Body Mass Index; OSAS: obstructive sleep apnea syndrome; AHI: apnea-hypopnea index. Age and BMI are reported as median [p25-p75], with p-value obtained from Mann-Whitney U test; effect size = rank-biserial correlation. Sex and clinical data are reported as number and percentage (%), with the p-value obtained from Chi-squared test; effect size = Phi coefficient.

**Whole night**

*Table S2a – Sleep macrostructure variables from PSG data in DIS with/without depressive symptoms.*

| Variable | DIS – dep  (n = 252) | DIS – no dep  (n =304) | Odds Ratio | Standard Error | p-value | CI 95% |
| --- | --- | --- | --- | --- | --- | --- |
| TIB (min) | 507 [485-539] | 507 [485-537] | 1.001 | 0.00151 | 0.571 | 0.998-1.00 |
| SPT (min) | 440 [394-475] | 443 [392-477] | 1.000 | 0.00121 | 0.790 | 0.997-1.00 |
| TST (min) | 369 [299-416] | 355 [305-406] | 1.000 | 0.00108 | 0.992 | 0.998-1.00 |
| Sleep efficiency (%) | 70.1 [58.2-79.5] | 68.0 [56.7-77.8] | 1.000 | 0.00584 | 0.939 | 0.989-1.01 |
| N1 (%) | 8.42 [5.50-13.4] | 9.02 [5.92-14.9] | 0.999 | 0.01174 | 0.903 | 0.976-1.02 |
| N2 (%) | 58.1 ± 10.6 | 59.3 ± 10.3 | 0.994 | 0.00863 | 0.517 | 0.978-1.01 |
| N3 (%) | 14.3 [5.97-21.1] | 11.5 [5.05-18.7] | 1.008 | 0.01010 | 0.413 | 0.989-1.03 |
| REM (%) | 17.5 [13.5-21.1] | 16.3 [12.2-21.2] | 1.001 | 0.01346 | 0.952 | 0.975-1.03 |
| Sleep latency (min) | 46.3 [22.4-86.1] | 44.3 [22.5-80.9] | 1.001 | 0.00169 | 0.513 | 0.998-1.00 |
| N2 latency (min) | 3 [1.5-6.5] | 3 [1-7.5] | 0.995 | 0.00675 | 0.500 | 0.982-1.01 |
| N3 latency (min) | 27.0 [15.5-56.3] | 36.3 [18.9-78.8] | 0.998 | 0.00151 | 0.173 | 0.995-1.00 |
| REM latency (min) | 97.8 [71.0-150] | 100 [67.5-157] | 1.001 | 0.00122 | 0.592 | 0.998-1.00 |
| WASO (min) | 53.3 [27-104] | 63.0 [32.8-115] | 0.999 | 0.00147 | 0.628 | 0.996-1.00 |
| oAHI (n/h) | 10 [3-24] | 15 [5-30] | 1.005 | 0.00536 | 0.296 | 0.994-1.02 |
| Desaturation index (n/h) | 8 [2-19.3] | 9 [3-25] | 1.008 | 0.00587 | 0.168 | 0.997-1.02 |
| Micro-arousal index (n/H) | 11.5 [7-20] | 13 [8-21] | 1.012 | 0.00772 | 0.137 | 0.996-1.03 |
| NREM microarousals (n/h) | 10.2 [5.5-17.4] | 10.9 [6.2-18.2] | 1.013 | 0.00800 | 0.118 | 0.997-1.03 |
| REM microarousals (n/h) | 1.71 [0.69-3.02] | 1.98 [0.93-3.44] | 0.996 | 0.04402 | 0.934 | 0.914-1.09 |
| LPM index (n/h) | 0.46 [0-4.96] | 0 [0-5.11] | 1.006 | 0.00627 | 0.367 | 0.993-1.02 |

TIB: time in bed; SPT: sleep period time; TST: total sleep time; AHI: apnea-hypopnea index; mA: micro-arousal. WASO: wake after sleep onset. Data are reported as median [p25-p75).

*Table S2b – Sleep macrostructure variables from PSG data in DIS with/without anxiety symptoms*

| Variable | DIS – anx  (n = 361) | DIS – no anx  (n = 194) | Odds Ratio | Standard Error | p-value | CI 95% |
| --- | --- | --- | --- | --- | --- | --- |
| TIB (min) | 511 [489-539] | 497 [481-534] | 1.004 | 0.00161 | **0.030** | 1.00-1.007 |
| SPT (min) | 444 [395-481] | 436 [392-474] | 1.001 | 0.00127 | 0.461 | 0.998-1.003 |
| TST (min) | 367 [303-417] | 351 [291-400] | 1.000 | 0.00113 | 0.906 | 0.998-1.002 |
| Sleep efficiency (%) | 69.9 [56.8-79.4] | 67.4 [57-76.6] | 0.998 | 0.00611 | 0.725 | 0.986-1.010 |
| N1 (%) | 8.17 [5.16-12.8] | 10.1 [6.73-16.3] | 0.987 | 0.01189 | 0.269 | 0.964-1.010 |
| N2 (%) | 58.4 ± 10 | 59.5 ± 11 | 0.999 | 0.00904 | 0.948 | 0.982-1.017 |
| N3 (%) | 13.5 [6.69-20.9] | 11.1 [3.24-17] | 1.010 | 0.01074 | 0.351 | 0.989-1.032 |
| REM (%) | 17 ± 6.42 | 16.2 ± 7.02 | 1.003 | 0.01408 | 0.859 | 0.975-1.031 |
| Sleep latency (min) | 46.5 [23.5-86.5] | 42 [21.3-77.4] | 1.003 | 0.00187 | 0.102 | 0.999-1.007 |
| N2 latency (min) | 3 [1-7] | 3 [1.5-7.5] | 0.996 | 0.00647 | 0.511 | 0.983-1.008 |
| N3 latency (min) | 28 [15.8-62.8] | 36.3 [20-77.5] | 1.001 | 0.00151 | 0.392 | 0.998-1.004 |
| REM latency (min) | 97 [69.8-151] | 103 [70-156] | 1.001 | 0.00129 | 0.378 | 0.999-1.004 |
| WASO (min) | 54.4 [27-106] | 65.8 [39.1-110] | 1.001 | 0.00152 | 0.554 | 0.998-1.004 |
| oAHI (n/h) | 10 [3-24] | 16 [6.25-30.8] | 1.004 | 0.00538 | 0.423 | 0.994-1.015 |
| Desaturation index (n/h) | 7 [2-19] | 10.5 [3-27] | 1.007 | 0.00595 | 0.215 | 0.996-1.019 |
| Micro-arousal index (n/h) | 11 [7-19] | 15 [8-23] | 0.997 | 0.00770 | 0.675 | 0.982-1.012 |
| NREM microarousals (n/h) | 9.92 [5.67-16.5] | 11.8 [6.26-20.5] | 0.999 | 0.00798 | 0.861 | 0.983-1.014 |
| REM microarousals (n/h) | 1.77 [0.73-3] | 2.04 [0.88-3.73] | 0.932 | 0.04581 | 0.124 | 0.852-1.020 |
| LPM index (n/h) | 0 [0-4.74] | 0.14 [0-6.55] | 1.001 | 0.00634 | 0.926 | 0.988-1.013 |

TIB: time in bed; SPT: sleep period time; TST: total sleep time; AHI: apnea-hypopnea index; mA: micro-arousal. WASO: wake after sleep onset.

Table S3a – Hypnodensity-based variables in DIS with/without depressive symptoms.

| Variable | DIS – dep  (n = 252) | DIS – no dep  (n = 304) | Odds Ratio | Standard Error | p_corr_ | CI 95% |
| --- | --- | --- | --- | --- | --- | --- |
| 5% sum W+N1 | 8 [3-20] | 8 [3-19] | 1.008 | 0.00432 | 0.156 | 0.999-1.02 |
| 10% sum W+N1 | 24.8 [10-56] | 26.0 [10.4-48] | 1.005 | 0.00268 | 0.133 | 1.000-1.01 |
| 30% sum W+N1 | 107 [69.4-171] | 110 [73.4-154] | 1.001 | 0.00134 | 0.483 | 0.998-1.00 |
| 50% sum W+N1 | 216 ± 80.8 | 213 ± 78.5 | 1.001 | 0.00111 | 0.483 | 0.999-1.00 |
| 5% sum N2+N3 | 28 [18.5-49.5] | 31.3 [19.9-59.1] | 0.995 | 0.00244 | 0.133 | 0.990-1.00 |
| 10% sum N2+N3 | 41.5 [29-70.6] | 47.5 [29-81.5] | 0.996 | 0.00203 | 0.133 | 0.992-1.00 |
| 30% sum N2+N3 | 114 [85.9-148] | 119 [85.9-155] | 0.999 | 0.00147 | 0.483 | 0.996-1.00 |
| 50% sum N2+N3 | 177 [139-226] | 181 [140-229] | 0.999 | 0.00125 | 0.483 | 0.997-1.00 |

Time to reach 5/10/30/50% of the cumulative sum between of the probability products of the set of stages.

Table S3b – Hypnodensity-based variables in DIS with/without anxiety symptoms.

| Variable | DIS - anxiety  (n = 361) | DIS - no anx  (n = 194) | Odds Ratio | Standard Error | p_corr_ | CI 95% |
| --- | --- | --- | --- | --- | --- | --- |
| 5% sum W+N1 | 7 [2.5-18] | 9.5 [4.5-20.4] | 1.007 | 0.00473 | 0.587 | 0.997-1.016 |
| 10% sum W+N1 | 22.5 [8.5-50.5] | 28.8 [13.1-51.3] | 1.004 | 0.00287 | 0.587 | 0.998-1.009 |
| 30% sum W+N1 | 107 [68.5-167] | 110 [78.1-150] | 1.003 | 0.00147 | 0.536 | 1.000-1.006 |
| 50% sum W+N1 | 215 ± 80.8 | 212 ± 78.4 | 1.001 | 0.00118 | 0.700 | 0.999-1.003 |
| 5% sum N2+N3 | 29.5 [18.5-54] | 31.3 [20-59.1] | 0.999 | 0.00231 | 0.752 | 0.995-1.004 |
| 10% sum N2+N3 | 45.5 [30-73] | 46.5 [28.5-78.8] | 0.999 | 0.00200 | 0.752 | 0.995-1.003 |
| 30% sum N2+N3 | 115 [85.5-153] | 118 [85.6-148] | 1.001 | 0.00153 | 0.752 | 0.998-1.004 |
| 50% sum N2+N3 | 178 [139-229] | 181 [142-227] | 1.001 | 0.00130 | 0.752 | 0.998-1.003 |

Time to reach 5/10/30/50% of the cumulative sum between of the probability products of the set of stages.

**Sleep Onset Period (SOP)**

Table S4 – Theta/Alpha ratio across SOP in patients with and without DIS.

| Variable | DIS  (n = 1328) | no DIS  (n = 1624) | Odds Ratio | Standard Error | p_corr_ | CI 95% |
| --- | --- | --- | --- | --- | --- | --- |
| Variance | 2.41 (1.22-5.28) | 2.44 (1.23-5.29) | 0.998 | 0.00150 | 0.353 | 0.996-1.001 |
| MSE (n) | 16 (3-43.3) | 15 (4-37) | 1.002 | 0.00051 | **0.012** | 1.001-1.003 |
| MSE index | 1.14 (0.22-2.25) | 1.14 (0.24-2.14) | 1.018 | 0.03306 | 0.585 | 0.954-1.086 |
| MSE duration (sec) | 67.5 (9-192) | 63 (12-180) | 1.000 | 0.00010 | **0.015** | 1.000-1.000 |
| AUC mean | 1.83 (1.11-2.82) | 1.88 (1.12-2.90) | 0.964 | 0.02363 | 0.174 | 0.920-1.009 |
| AUC index | 0.09 (0.03-0.20) | 0.10 (0.03-0.22) | 0.637 | 0.23284 | 0.106 | 0.404-1.005 |

MSE: micro-sleep event; MSE (n): number of micro-sleep events; MSE index: ratio between number of micro-sleep events and sleep onset period length; MSE duration: sum of duration of each micro-sleep event; AUC: area under the curve

Table S5 – iVS across SOP in patients with and without DIS.

| Variable | DIS  (n = 1328) | no DIS  (n = 1624) | Odds Ratio | Standard Error | p_corr_ | CI 95% |
| --- | --- | --- | --- | --- | --- | --- |
| iVS variance | 1364 (1180-1552) | 1384 (1189-1575) | 1.000 | 0.00013 | 0.378 | 1.000-1.000 |
| MSE (n) | 29 (15-62.3) | 28 (15-57) | 1.002 | 0.00050 | **0.005** | 1.001-1.003 |
| MSE index | 1.86 (1.34-2.40) | 1.83 (1.33-2.33) | 1.027 | 0.05062 | 0.664 | 0.930-1.134 |
| MSE duration (sec) | 165 (84-342) | 159 (84-306) | 1.000 | 0.00009 | **0.014** | 1.000-1.000 |
| AUC mean | 1.81 (1.24-2.56) | 1.87 (1.23-2.52) | 0.996 | 0.03718 | 0.911 | 0.926-1.071 |
| AUC index | 0.109 (0.04-0.21) | 0.111 (0.04-0.21) | 0.698 | 0.27306 | 0.315 | 0.409-1.193 |

The corrected p-value refers to the logistic regression analysis. MSE: micro-sleep events; AUC: area under the curve. All indexes refer to number/time.

Table S6 – Hypnodensity-derived stage probability during SOP in patients with and without DIS.

| Variable | DIS  (n=1328) | no DIS  (n=1624) | Odds Ratio | Standard Error | p_corr_ | CI 95% |
| --- | --- | --- | --- | --- | --- | --- |
| Probability W (%) | 63.2 (48.6-77.1) | 64.3 (49.6-78.0) | 0.996 | 0.00198 | 0.276 | 0.993-1.000 |
| Probability N1 (%) | 15.3 (9.26-21.9) | 15.6 (9.66-22.9) | 0.995 | 0.00419 | 0.416 | 0.987-1.003 |
| Probability N2 (%) | 10.7 (5.07-18.6) | 11.4 (5.68-19.2) | 1.001 | 0.00363 | 0.801 | 0.994-1.008 |
| Probability N3 (%) | 0.08 (0.02-0.31) | 0.07 (0.02-0.30) | 1.017 | 0.01636 | 0.416 | 0.985-1.050 |

Table S7a– Sleep Onset Period in DIS with/without depressive symptoms.

| Variable | DIS – dep  (n = 252) | DIS – no dep  (n = 304) | Odds Ratio | Standard Error | p_corr_ | CI 95% |
| --- | --- | --- | --- | --- | --- | --- |
| SOP length (min) | 11.3 [7-22.6] | 12.5 [7-34.3] | 0.998 | 0.00209 | 0.654 | 0.994-1.00 |
| W duration (min) | 4.5 [3-10.6] | 5.75 [3.5-16.6] | 0.996 | 0.00299 | 0.548 | 0.990-1.00 |
| N1 duration (min) | 3.5 [2-6.5] | 3.5 [1.5-7] | 1.003 | 0.01030 | 0.933 | 0.983-1.02 |
| N2 duration (min) | 2 [1-5] | 2 [1-5] | 1.001 | 0.01636 | 0.933 | 0.970-1.03 |

Table S7b– Sleep Onset Period in DIS with/without anxiety symptoms.

| Variable | DIS – anx  (n = 361) | DIS – no anx  (n = 194) | Odds Ratio | Standard Error | p_corr_ | CI 95% |
| --- | --- | --- | --- | --- | --- | --- |
| SOP length (min) | 11 [7-24] | 14.5 [7.5-35.8] | 0.998 | 0.00199 | 0.478 | 0.994-1.002 |
| W duration (min) | 5 [3-12] | 6.5 [3.5-17.9] | 0.998 | 0.00260 | 0.478 | 0.993-1.003 |
| N1 duration (min) | 3.5 [2-6.5] | 4 [2-8] | 0.988 | 0.01128 | 0.478 | 0.967-1.010 |
| N2 duration (min) | 2 [1-4.5] | 2 [1-6] | 0.988 | 0.01642 | 0.478 | 0.957-1.021 |

Table S8a – Theta/Alpha ratio across SOP in DIS with/without depression.

| Variable | DIS – dep  (n = 252) | DIS – no dep  (n = 304) | Odds Ratio | Standard Error | p_corr_ | CI 95% |
| --- | --- | --- | --- | --- | --- | --- |
| Variance | 2.91 [1.27-6.26] | 2.33 [1.04-4.45] | 1.001 | 0.00297 | 0.650 | 0.996-1.01 |
| MSE (n) | 17 [4-39.5] | 16.5 [4-49] | 0.999 | 0.00100 | 0.650 | 0.997-1.00 |
| MSE index | 1.26 [0.23-2.31] | 1.20 [0.21-2.22] | 1.045 | 0.07686 | 0.650 | 0.899-1.21 |
| MSE duration (sec) | 76.5 [12-198] | 73.5 [12-222] | 1.000 | 0.00020 | 0.650 | 1.000-1.00 |
| AUC mean | 2.06 [1.28-3.20] | 1.80 [1.24-2.60] | 1.031 | 0.04607 | 0.650 | 0.942-1.13 |
| AUC index | 0.11 [0.03-0.24] | 0.08 [0.02-0.19] | 1.897 | 0.54252 | 0.650 | 0.655-5.49 |

MSE: micro-sleep event; MSE (n): number of micro-sleep events; MSE index: ratio between number of micro-sleep events and sleep onset period length; MSE duration: sum of duration of each micro-sleep event; AUC: area under the curve

Table S8b– iVS across SOP in DIS with/without depressive symptoms.

| Variable | DIS – dep  (n = 252) | DIS – no dep  (n = 304) | Odds Ratio | Standard Error | p_corr_ | CI 95% |
| --- | --- | --- | --- | --- | --- | --- |
| iVS variance | 1437 [1223-1626] | 1390 [1201-1546] | 1.000 | 0.00029 | 0.258 | 0.999-1.00 |
| iVS mean | 45.9 [36.5-54.3] | 48.4 [39.8-59.8] | 0.979 | 0.00648 | **0.005** | 0.966-0.991 |
| MSE (n) | 27.5 [13-51.3] | 30 [16-79.3] | 0.998 | 0.00114 | 0.197 | 0.996-1.00 |
| MSE index | 1.82 ± 0.75 | 1.82 ± 0.74 | 1.015 | 0.11694 | 0.989 | 0.807-1.28 |
| MSE duration (sec) | 156 [83.3-286] | 168 [87-402] | 1.000 | 0.00021 | 0.322 | 0.999-1.00 |
| AUC mean | 1.81 [1.21-2.56] | 1.76 [1.27-2.47] | 1.039 | 0.08725 | 0.822 | 0.876-1.23 |
| AUC index | 0.11 [0.05-0.198] | 0.09 [0.03-0.202] | 0.992 | 0.64004 | 0.989 | 0.283-3.48 |

MSE: micro-sleep event; MSE (n): number of micro-sleep events; MSE index: ratio between number of micro-sleep events and sleep onset period length; MSE duration: sum of duration of each micro-sleep event; AUC: area under the curve

Table S9a – Theta/Alpha ratio across SOP in DIS with/without anxiety symptoms.

| Variable | DIS – anx  (n = 361) | DIS – no anx  (n = 194) | Odds Ratio | Standard Error | p_corr_ | CI 95% |
| --- | --- | --- | --- | --- | --- | --- |
| Variance | 2.68 [1.21-5.87] | 2.14 [1.05-4.42] | 1.005 | 0.00535 | 0.661 | 0.995-1.015 |
| MSE (n) | 16 [4-42] | 18.5 [4-55.5] | 0.999 | 0.00089 | 0.661 | 0.998-1.001 |
| MSE index | 1.27 [0.23-2.25] | 1.14 [0.19-2.23] | 1.007 | 0.07981 | 0.928 | 0.861-1.178 |
| MSE duration (sec) | 75 [12-192] | 78 [12-252] | 1.000 | 0.00018 | 0.661 | 1.000-1.000 |
| AUC mean | 1.96 [1.20-2.99] | 1.81 [1.27-2.66] | 0.961 | 0.04725 | 0.661 | 0.876-1.054 |
| AUC index | 0.09 [0.03-0.23] | 0.08 [0.02-0.17] | 2.168 | 0.61076 | 0.661 | 0.655-7.177 |

MSE: micro-sleep event; MSE (n): number of micro-sleep events; MSE index: ratio between number of micro-sleep events and sleep onset period length; MSE duration: sum of duration of each micro-sleep event; AUC: area under the curve.

*Table S9b – iVS across SOP in DIS with/without anxiety symptoms.*

| Variable | DIS – anx  (n = 361) | DIS – no anx  (n = 194) | Odds Ratio | Standard Error | p_corr_ | CI 95% |
| --- | --- | --- | --- | --- | --- | --- |
| iVS variance | 1401 [1196-1580] | 1399 [1238-1553] | 1.000 | 0.00031 | 0.691 | 0.999-1.000 |
| iVS mean | 46.2 [37.7-57.9] | 49.3 [40-58.9] | 0.991 | 0.00642 | 0.582 | 0.979-1.004 |
| %epochs 0-20 | 33.7 ± 14 | 31.2 ± 13.4 | 1.010 | 0.00675 | 0.582 | 0.996-1.023 |
| %epochs 20-66 | 24.3 [18.5-31.4] | 24.4 [19-31] | 1.001 | 0.01003 | 0.916 | 0.982-1.021 |
| %epochs 66-100 | 39.5 [29.1-52.4] | 41.6 [31.9-52.7] | 0.993 | 0.00548 | 0.582 | 0.983-1.004 |
| MSE (n) | 26 [14-55] | 33.5 [16-81.3] | 0.999 | 0.00105 | 0.691 | 0.997-1.001 |
| MSE index | 1.77 [1.30-2.27] | 1.87 [1.32-2.44] | 0.919 | 0.12332 | 0.691 | 0.721-1.170 |
| MSE duration (sec) | 153 [84-294] | 177 [87-402] | 1.000 | 0.00020 | 0.767 | 1.000-1.000 |
| AUC mean | 1.78 [1.24-2.55] | 1.77 [1.26-2.45] | 1.091 | 0.09389 | 0.691 | 0.908-1.311 |
| AUC index | 0.11 [0.05-0.20] | 0.09 [0.03-0.19] | 2.322 | 0.70553 | 0.582 | 0.582-9.255 |

MSE: micro-sleep event; MSE (n): number of micro-sleep events; MSE index: ratio between number of micro-sleep events and sleep onset period length; MSE duration: sum of duration of each micro-sleep event; AUC: area under the curve

**AASM Pre-sleep Wakefulness**

Table S10 –iVS across pre-sleep wakefulness in patients with and without DIS.

| Variable | DIS  (n = 1328) | no DIS  (n = 1624) | Odds Ratio | Standard Error | p_corr_ | CI 95% |
| --- | --- | --- | --- | --- | --- | --- |
| iVS variance | 527 (356-694) | 533 (373-694) | 1.000 | 0.00017 | 0.271 | 0.999-1.000 |
| MSE (n) | 4 (1-7) | 3 (1-6) | 1.028 | 0.01112 | **0.042** | 1.006-1.051 |
| MSE index | 1.33 (0.33-2.33) | 1 (0.33-2) | 1.088 | 0.03336 | **0.042** | 1.019-1.161 |
| MSE duration (sec) | 12 (3-24) | 12 (3-21) | 1.006 | 0.00262 | **0.042** | 1.001-1.011 |
| AUC mean | 0.295 (0.09-0.59) | 0.27 (0.07-0.51) | 1.185 | 0.07946 | **0.048** | 1.015-1.385 |
| AUC index | 0.098 (0.03-0.19) | 0.089 (0.02-0.17) | 1.666 | 0.23838 | **0.048** | 1.044-2.658 |

The corrected p-value refers to the logistic regression analysis. MSE: micro-sleep events; AUC: area under the curve. All indexes refer to number/time.

Table S11 – Theta/Alpha Ratio across pre-sleep wakefulness in patients with and without DIS.

| Variable | DIS  (n = 1328) | no DIS  (n = 1624) | Odds Ratio | Standard Error | p_corr_ | CI 95% |
| --- | --- | --- | --- | --- | --- | --- |
| Variance | 1.09 (0.53-2.10) | 1.06 (0.55-2.03) | 0.997 | 0.00676 | 0.754 | 0.984-1.010 |
| MSE (n) | 1 (0-5) | 1 (0-5) | 1.008 | 0.01103 | 0.754 | 0.987-1.030 |
| MSE index | 0.33 (0-1.67) | 0.33 (0-1.67) | 1.025 | 0.03309 | 0.754 | 0.961-1.094 |
| MSE duration (sec) | 3 (0-18) | 3 (0-18) | 1.002 | 0.00233 | 0.754 | 0.997-1.006 |
| AUC mean | 0.74 (0-1.84) | 0.68 (0-1.79) | 1.007 | 0.02132 | 0.754 | 0.966-1.050 |
| AUC index | 0.25 (0-0.61) | 0.23 (0-0.59) | 1.020 | 0.06395 | 0.754 | 0.900-1.156 |

The corrected p-value refers to the logistic regression analysis. MSE: micro-sleep event; AUC: area under the curve. Variance refers to the variance of theta/alpha ratio across the 3 minutes before sleep. All indexes refer to number/time.

Table S12a – Theta/Alpha Ratio across pre-sleep wakefulness in groups with DIS with/without depressive symptoms.

| Variable | DIS – dep  (n = 252) | DIS – no dep  (n = 304) | Odds Ratio | Standard Error | p_corr_ | CI 95% |
| --- | --- | --- | --- | --- | --- | --- |
| Variance | 1.19 [0.53-2.27] | 1.04 [0.52-1.96] | 1.007 | 0.01419 | 0.643 | 0.979-1.03 |
| MSE (n) | 1 [0-4] | 1 [0-5] | 1.016 | 0.02561 | 0.643 | 0.966-1.07 |
| MSE index | 0.33 [0-1.33] | 0.33 [0-1.67] | 1.048 | 0.07683 | 0.643 | 0.901-1.22 |
| MSE duration (sec) | 6 [0-18] | 3 [0-15] | 1.005 | 0.00536 | 0.643 | 0.995-1.02 |
| AUC mean | 0.84 [0-1.88] | 0.68 [0-1.85] | 1.041 | 0.04776 | 0.643 | 0.948-1.14 |
| AUC index | 0.28 [0-0.63] | 0.23 [0-0.62] | 1.127 | 0.14327 | 0.643 | 0.851-1.49 |

MSE: micro-sleep event; MSE (n): number of micro-sleep events; MSE index: ratio between number of micro-sleep events and sleep onset period length; MSE duration: sum of duration of each micro-sleep event; AUC: area under the curve

Table S12b – iVS across pre-sleep wakefulness in groups with DIS with/without depressive symptoms.

| Variable | DIS – dep  (n = 252) | DIS – no dep  (n = 304) | Odds Ratio | Standard Error | p_corr_ | CI 95% |
| --- | --- | --- | --- | --- | --- | --- |
| iVS variance | 535 ± 237 | 528 ± 220 | 1.000 | 0.00039 | 0.797 | 0.999-1.00 |
| iVS mean | 79.5 [71.3-87.8] | 80.8 [72.5-87.5] | 0.992 | 0.00825 | 0.453 | 0.976-1.01 |
| %epochs 0-20 | 0 [0-3.33] | 0 [0-1.67] | 1.059 | 0.03230 | 0.453 | 0.994-1.13 |
| %epochs 20-66 | 23.3 [11.7-36.7] | 21.7 [11.7-33.8] | 1.004 | 0.00569 | 0.488 | 0.993-1.02 |
| %epochs 66-100 | 75.8 [60-88.3] | 76.7 [63.3-86.7] | 0.995 | 0.00519 | 0.453 | 0.985-1.00 |
| MSE (n) | 3 [1-6] | 4 [1-6] | 0.972 | 0.02719 | 0.453 | 0.921-1.02 |
| MSE index | 1 [0.33-2] | 1.33 [0.33-2] | 0.917 | 0.08158 | 0.453 | 0.782-1.08 |
| MSE duration (sec) | 12 [3-21] | 12 [3-24] | 0.995 | 0.00636 | 0.488 | 0.983-1.01 |
| AUC mean | 0.26 [0.09-0.53] | 0.33 [0.11-0.59] | 0.805 | 0.19099 | 0.453 | 0.554-1.17 |
| AUC index | 0.09 [0.03-0.17] | 0.11 [0.04-0.19] | 0.522 | 0.57298 | 0.453 | 0.170-1.61 |

MSE: micro-sleep event; MSE (n): number of micro-sleep events; MSE index: ratio between number of micro-sleep events and sleep onset period length; MSE duration: sum of duration of each micro-sleep event; AUC: area under the curve

Table S13a – Theta/Alpha Ratio across pre-sleep wakefulness in DIS with/without anxiety symptoms.

| Variable | DIS – anx  (n = 361) | DIS – no anx  (n = 194) | Odds Ratio | Standard Error | p_corr_ | CI 95% |
| --- | --- | --- | --- | --- | --- | --- |
| Variance | 1.01 [0.49-2.13] | 1.22 [0.57-2.09] | 1.014 | 0.01851 | 0.988 | 0.978-1.052 |
| MSE (n) | 1 [0-4] | 2 [0-4] | 1.000 | 0.02637 | 0.988 | 0.949-1.053 |
| MSE index | 0.33 [0-1.33] | 0.67 [0-1.33] | 0.999 | 0.07910 | 0.988 | 0.855-1.166 |
| MSE duration (sec) | 3 [0-18] | 6 [0-15] | 0.998 | 0.00548 | 0.988 | 0.987-1.009 |
| AUC mean | 0.76 [0-1.88] | 0.70 [0-1.93] | 0.999 | 0.05090 | 0.988 | 0.904-1.104 |
| AUC index | 0.25 [0-0.63] | 0.23 [0-0.64] | 0.998 | 0.15271 | 0.988 | 0.740-1.346 |

MSE: micro-sleep event; MSE (n): number of micro-sleep events; MSE index: ratio between number of micro-sleep events and sleep onset period length; MSE duration: sum of duration of each micro-sleep event; AUC: area under the curve

Table S13b – iVS across pre-sleep wakefulness in DIS with/without anxiety symptoms.

| Variable | DIS – anx  (n = 361) | DIS – no anx  (n = 194) | Odds Ratio | Standard Error | p_corr_ | CI 95% |
| --- | --- | --- | --- | --- | --- | --- |
| iVS variance | 529 [359-684] | 523 [358-685] | 1.000 | 0.00041 | 0.898 | 0.999-1.001 |
| iVS mean | 80.1 [71.7-87.6] | 80.6 [72.5-86.9] | 0.997 | 0.00863 | 0.898 | 0.981-1.014 |
| %epochs 0-20 | 0 [0-3.33] | 0 [0-1.67] | 1.053 | 0.03669 | 0.898 | 0.980-1.132 |
| %epochs 20-66 | 23.3 [11.7-36.7] | 22.5 [13.3-35] | 1.001 | 0.00594 | 0.898 | 0.989-1.012 |
| %epochs 66-100 | 76.7 [60-88.3] | 76.7 [63.3-86.3] | 0.998 | 0.00543 | 0.898 | 0.988-1.009 |
| MSE (n) | 3 [1-6] | 4 [1-6] | 0.985 | 0.02778 | 0.898 | 0.933-1.040 |
| MSE index | 1 [0.33-2] | 1.33 [0.33-2] | 0.956 | 0.08333 | 0.898 | 0.812-1.125 |
| MSE duration (sec) | 12 [3-21] | 12 [3-24] | 0.998 | 0.00652 | 0.898 | 0.985-1.010 |
| AUC mean | 0.28 [0.09-0.56] | 0.29 [0.11-0.60] | 0.836 | 0.19253 | 0.898 | 0.574-1.220 |
| AUC index | 0.09 [0.03-0.19] | 0.09 [0.03-0.20] | 0.585 | 0.57760 | 0.898 | 0.189-1.815 |

MSE: micro-sleep event; MSE (n): number of micro-sleep events; MSE index: ratio between number of micro-sleep events and sleep onset period length; MSE duration: sum of duration of each micro-sleep event; AUC: area under the curve.

**Early consolidated sleep**

Table S14 – Theta/Alpha Ratio across first 10 minutes of consolidated sleep in patients with and without DIS.

| Variable | DIS  (n = 1328) | no DIS  (n = 1624) | Odds Ratio | Standard Error | p_corr_ | CI 95% |
| --- | --- | --- | --- | --- | --- | --- |
| Variance | 2.75 (1.25-6.11) | 2.90 (1.34-6.38) | 1.000 | 0.00106 | 0.884 | 0.997-1.002 |
| MSE (n) | 12 (2-32) | 12.5 (2-32) | 1.001 | 0.00239 | 0.884 | 0.996-1.005 |
| MSE index | 1.2 (0.2-3.2) | 1.25 (0.2-3.2) | 1.008 | 0.02385 | 0.884 | 0.962-1.056 |
| MSE duration (sec) | 48 (6-150) | 51 (6-153) | 1.000 | 0.00044 | 0.961 | 0.999-1.001 |
| AUC mean | 1.74 (0.76-3.01) | 1.76 (0.80-3.05) | 0.993 | 0.01427 | 0.884 | 0.966-1.021 |
| AUC index | 0.17 (0.07-0.30) | 0.18 (0.08-0.31) | 0.933 | 0.14273 | 0.884 | 0.705-1.234 |

The corrected p-value refers to the logistic regression analysis. MSE: micro-sleep event; MSE (n): number of micro-sleep events; MSE index: ratio between number of micro-sleep events and time; MSE duration: sum of duration of each micro-sleep event, expressed in seconds; AUC: area under the curve; AUC index: ratio between AUC and time.

Table S15 – iVS across first 10 minutes of consolidated sleep in patients with and without DIS.

| Variable | DIS  (n = 1328) | no DIS  (n = 1624) | Odds Ratio | Standard Error | p_corr_ | CI 95% |
| --- | --- | --- | --- | --- | --- | --- |
| iVS variance | 491 (315-682) | 441 (282-628) | 1.001 | 0.00016 | **0.002** | 1.000-1.001 |
| MSE (n) | 8 (3-19) | 8 (3-17) | 1.005 | 0.00324 | 0.161 | 0.999-1.012 |
| MSE index | 0.8 (0.3-1.9) | 0.8 (0.3-1.7) | 1.053 | 0.03239 | 0.161 | 0.988-1.122 |
| MSE duration (sec) | 60 (18-138) | 54 (18-126) | 1.001 | 0.00050 | 0.254 | 1.000-1.002 |
| AUC mean | 2.84 (1.37-4.47) | 2.69 (1.39-4.43) | 0.999 | 0.01399 | 0.929 | 0.972-1.027 |
| AUC index | 0.28 (0.14-0.45) | 0.27 (0.14-0.44) | 0.988 | 0.13985 | 0.929 | 0.751-1.299 |

The corrected p-value refers to the logistic regression analysis. MSE: micro-sleep events; AUC: area under the curve. All indexes refer to number/time.

Table S16a – Theta/Alpha Ratio across first 10 minutes of consolidated sleep in DIS with/without depressive symptoms.

| Variable | DIS – dep  (n = 252) | DIS – no dep  (n = 304) | Odds Ratio | Standard Error | p_corr_ | CI 95% |
| --- | --- | --- | --- | --- | --- | --- |
| Variance | 3.09 [1.30-7.45] | 2.81 [1.32-6.22] | 1.000 | 0.00161 | 0.884 | 0.996-1.00 |
| MSE (n) | 10 [2-30.3] | 14.5 [2-34] | 0.996 | 0.00541 | 0.630 | 0.985-1.01 |
| MSE index | 1 [0.2-3.02] | 1.45 [0.2-3.4] | 0.957 | 0.05406 | 0.630 | 0.861-1.06 |
| MSE duration (sec) | 40.5 [6-148] | 57 [6-154] | 1.000 | 0.00099 | 0.884 | 0.998-1.00 |
| AUC mean | 1.91 [0.82-3.33] | 1.80 [0.97-3.06] | 1.052 | 0.03205 | 0.342 | 0.988-1.12 |
| AUC index | 0.19 [0.08-0.33] | 0.18 [0.09-0.31] | 1.660 | 0.32050 | 0.342 | 0.886-3.11 |

Table S16b – iVS across first 10 minutes of consolidated sleep in DIS with/without depressive symptoms.

| Variable | DIS – dep  (n = 252) | DIS – no dep  (n = 304) | Odds Ratio | Standard Error | p_corr_ | CI 95% |
| --- | --- | --- | --- | --- | --- | --- |
| iVS variance | 433 [286-660] | 460 [310-641] | 1.000 | 0.00036 | 0.955 | 0.999-1.00 |
| iVS mean | 10.6 [7.12-16.2] | 11 [7.30-16.2] | 1.001 | 0.01181 | 0.955 | 0.978-1.02 |
| %epochs 0-20 | 78.0 [69.5-85] | 78.5 [67.9-85.5] | 0.999 | 0.00635 | 0.955 | 0.986-1.01 |
| %epochs 20-66 | 16.6 [11.4-23] | 16 [10.9-24] | 1.003 | 0.00276 | 0.955 | 0.986-1.02 |
| %epochs 66-100 | 4.5 [2.5-8.5] | 5 [3-8] | 0.999 | 0.01627 | 0.955 | 0.968-1.03 |
| MSE (n) | 8 [3-18.3] | 7 [2-14] | 1.013 | 0.00782 | 0.340 | 0.998-1.03 |
| MSE index | 0.8 [0.3-1.83] | 0.7 [0.2-1.4] | 1.139 | 0.07820 | 0.340 | 0.977-1.33 |
| MSE duration (sec) | 57 [18-132] | 51 [18-105] | 1.002 | 0.00120 | 0.430 | 1.000-1.00 |
| AUC mean | 2.96 [1.45-4.25] | 2.67 [1.35-4.48] | 1.022 | 0.03061 | 0.952 | 0.963-1.09 |
| AUC index | 0.29 [0.14-0.42] | 0.27 [0.13-0.45] | 1.244 | 0.30614 | 0.952 | 0.683-2.27 |

MSE: micro-sleep event; MSE (n): number of micro-sleep events; MSE index: ratio between number of micro-sleep events and sleep onset period length; MSE duration: sum of duration of each micro-sleep event; AUC: area under the curve

Table S17a – Theta/Alpha Ratio across first 10 minutes of consolidated sleep in DIS with/without anxiety symptoms.

| Variable | DIS – anx  (n = 361) | DIS – no anx  (n = 194) | Odds Ratio | Standard Error | p_corr_ | CI 95% |
| --- | --- | --- | --- | --- | --- | --- |
| Variance | 3.18 [1.40-6.78] | 2.49 [1.27-6.18] | 1.002 | 0.00242 | 0.716 | 0.997-1.007 |
| MSE (n) | 14 [2-32] | 12 [2-31.8] | 1.002 | 0.00566 | 0.716 | 0.991-1.013 |
| MSE index | 1.4 [0.2-3.20] | 1.2 [0.2-3.18] | 1.021 | 0.05658 | 0.716 | 0.914-1.141 |
| MSE duration (sec) | 57 [6-153] | 43.5 [6-153] | 1.001 | 0.00106 | 0.716 | 0.999-1.003 |
| AUC mean | 1.87 [0.84-3.27] | 1.81 [0.95-2.93] | 1.064 | 0.03750 | 0.297 | 0.988-1.145 |
| AUC index | 0.19 [0.08-0.33] | 0.18 [0.09-0.29] | 1.856 | 0.37496 | 0.297 | 0.890-3.870 |

MSE: micro-sleep event; MSE (n): number of micro-sleep events; MSE index: ratio between number of micro-sleep events and sleep onset period length; MSE duration: sum of duration of each micro-sleep event; AUC: area under the curve

Table S17b – iVS across first 10 minutes of consolidated sleep in DIS with/without anxiety symptoms.

| Variable | DIS – anx  (n = 361) | DIS – no anx  (n = 194) | Odds Ratio | Standard Error | p_corr_ | CI 95% |
| --- | --- | --- | --- | --- | --- | --- |
| iVS variance | 452 [309-641] | 457 [282-651] | 1.000 | 0.00038 | 0.766 | 0.999-1.001 |
| iVS mean | 11.2 [7.66-16] | 10.2 [6.79-16.9] | 1.009 | 0.01235 | 0.615 | 0.985-1.034 |
| %epochs 0-20 | 78 [69-84.5] | 79.8 [67-86.5] | 0.992 | 0.00676 | 0.615 | 0.979-1.006 |
| %epochs 20-66 | 16.5 [12-23.5] | 15.5 [10-24.5] | 1.013 | 0.00946 | 0.615 | 0.994-1.031 |
| %epochs 66-100 | 4.5 [2.5-8] | 5 [2.5-8] | 1.008 | 0.01694 | 0.689 | 0.975-1.042 |
| MSE (n) | 8 [3-18] | 7 [2-14] | 1.006 | 0.00820 | 0.615 | 0.990-1.022 |
| MSE index | 0.8 [0.3-1.8] | 0.7 [0.2-1.4] | 1.058 | 0.08199 | 0.615 | 0.901-1.242 |
| MSE duration (sec) | 57 [18-126] | 43.5 [15-105] | 1.001 | 0.00126 | 0.615 | 0.999-1.004 |
| AUC mean | 2.89 [1.48-4.39] | 2.59 [1.14-4.46] | 1.041 | 0.03363 | 0.615 | 0.975-1.112 |
| AUC index | 0.29 [0.15-0.44] | 0.26 [0.11-0.45] | 1.494 | 0.33633 | 0.615 | 0.773-2.888 |

MSE: micro-sleep event; MSE (n): number of micro-sleep events; MSE index: ratio between number of micro-sleep events and sleep onset period length; MSE duration: sum of duration of each micro-sleep event; AUC: area under the curve
